# Design of a Soluble Multivalent Notch Agonist

**DOI:** 10.1101/2024.08.07.607099

**Authors:** Rubul Mout, Ran Jing, Emily D Egan, Helen Eisenach, Mohamad Ali Toufic Najia, Luca Hensch, Trevor Bingham, Martin A. Kononov, Rajesh Gunage, Yunliang Zhao, Natasha I Edman, Thorsten M Schlaeger, Leonard I Zon, Trista E North, Urban Lendahl, David Baker, Stephen C Blacklow, George Q Daley

## Abstract

Designed protein agonists can enhance the efficiency of endogenous signaling pathways, and provide a powerful means to control cellular functions and develop disease therapeutics. Designing a soluble cytokine-like agonist for Notch signaling, an evolutionarily conserved pathway that regulates cell fate in embryonic and adult development, is especially challenging because Notch receptor activation requires a mechanical force that is typically mediated by cell-associated transmembrane ligands at sites of cell-cell contact. Moreover, free soluble Notch ligand is signal inhibitory. Here, we exploit computationally designed protein oligomers with precise geometries and valencies to generate cytokine-like, ‘protein only,’ multivalent soluble Notch agonists. These tools promote cell-cell contact, cluster Notch proteins in synapses at the cell surface, and activate Notch signaling in reporter cell lines and cells expressing endogenous receptors. We demonstrate the utility of these soluble Notch agonists in T cell differentiation from cord blood (CB) and human induced pluripotent stem cells (iPSCs), and in bioreactor production of T cells in liquid suspension. Soluble multivalent Notch agonists can be applied broadly to *in vitro* cellular differentiation methods to generate clinical cell products and to develop immunotherapies.

## Introduction

Notch signaling is an evolutionarily conserved pathway and a key developmental fate specifier in numerous cell types, including immune cells, neurons, vascular endothelial cells, and cardiomyocytes, among others^1^. Consequently, *in vitro* generation of many cell types from progenitor cells requires activating Notch signaling. Notch activation occurs through cell-cell interaction between sender cells, which present a repertoire of transmembrane Notch ligands (*i.e.*, Delta-like ligand (DLL1,3,4) and Jagged(1,2)), and receiver cells, which bind the ligand through Notch receptors, Notch(1-4)^2^. Receptor activation requires the delivery of mechanical force by the bound ligand to induce Notch proteolysis, which liberates the intracellular domain of the Notch receptor (Notch intracellular domain, NICD) to access the nucleus and induce transcription of NICD-responsive genes^1–3^. Due to its mechanical force requirement, Notch activation in cell culture requires the presentation of a natural transmembrane Notch ligand by sender cells, or by an immobilized ligand ectodomain on microbeads or on a tissue-culture dish. Although there are recent claims that Fc-DLL1 or the DNA origami-based presentation of a Notch ligand can induce a Notch signaling response, these activities have a very low signal-to-noise ratio, the mechanism is not understood, and their ability to induce cellular differentiation *in vitro* was not explored^4,5^. These recent data are also inconsistent with the long-standing, larger body of evidence showing that soluble Notch ligands inhibit signaling unless immobilized^6^. A DLL4-Fc fusion protein, for example, is a well-validated inhibitory decoy that blocks Notch signaling *in vivo* with efficacy comparable to that of an inhibitory antibody^7^.

Notch signaling is involved at various stages of hematopoietic development and differentiation, including specification of definitive hematopoietic stem cells (HSC), and maturation of T cells and their peripheral effector function^8,9^. A soluble Notch agonist could inform our understanding of Notch signaling *in vivo,* and contribute to the development of new immunotherapies including anti-tumor immunity and vaccine development. Of particular relevance, T cells derived *in vitro* from human induced pluripotent stem cells (iPSCs) have gained attention for their potential as “off-the-shelf” adoptive cell therapy in immune deficiency, viral infection, autoimmunity, and cancer^10,11^. More efficient manufacture of T cell products would facilitate clinical applications.

While human iPSCs provide an inexhaustible source of starting cell material for *in vitro* production of T cells, large-scale manufacturing of cell products still faces significant hurdles. Notch signaling is essential for T cell generation from iPSCs. In previous reports, Delta-like ligand 4 (DLL4) was presented to T cells by expression on adherent stromal cells^12,13^, coating of microbeads^14^, or immobilization on a tissue-culture plate^15–17^. Thus, the scalable manufacture of T cell products is hampered by the two-dimensional (2D) presentation of Notch ligands. We sought to develop a soluble Notch ligand that could induce receptor signaling in liquid suspension, thereby facilitating the manufacturing of T cells in a bioreactor.

To fabricate such a soluble Notch agonist, we took advantage of recent developments in computational protein design^18,19^, which specifies secondary, tertiary, and quaternary structures with atomically precise geometries and valencies, and which has been used in many types of signal activation^20–22^. Using the Rosetta protein design tool, we built multivalent protein oligomers with distinct geometries and numerical valencies (*i.e.*, 2, 3, 5, 6, 8, 60, and 120)^23–25^, coupled the Notch ligand DLL4 to the oligomeric scaffolds, and tested their capacity for receptor activation in reporter cell lines and cells with endogenous Notch signaling. Here, we describe the development of a soluble Notch-receptor agonist and demonstrate the production of T cells in a 3D bioreactor culture system, which vastly improves the yield and cost efficiency of *in vitro* T cell differentiation.

## Results

### Design of Soluble Notch agonists

We hypothesized that the *trans*-binding of multivalent soluble DLL4 ligands would bridge cells through ligand-receptor interactions, exert a physical force on the receptor, and activate Notch signaling (**Figure 1A**). Furthermore, we surmised that a multivalent ligand might amplify signaling by facilitating receptor clustering. We employed a previously described library of computationally designed proteins featuring diverse valencies, sizes, and geometries as scaffolds to engineer multivalent DLL4 complexes^23,24^. Covalent crosslinking of these oligomeric scaffolds to the N-terminus of the extracellular domain of human DLL4 was conducted using SpyLigation, in which a covalent isopeptide bond is formed between SpyCatcher (**SC**, 88 amino acids, 9.5 kDa) and SpyTag (**ST**, 13 amino acids) species^26,27^ (**Figure 1B**). All oligomeric scaffolds were fused to SC with a flexible glycine-serine (GS) linker and expressed and purified in *E. coli*. The DLL4 protein was fused to ST with a GS linker and expressed and purified in mammalian Expi293F cells. Multivalent building blocks were selected from a previously designed homodimer^23^ (C2), homotrimer^24^ (C3), helical bundle homotrimer^28^ (C3-HB), homopentamer^24^ (C5), homohexamer^25^ (C6), hexavalent dihedral oligomer^29^ (D3), homo-octamer^25^ (C8), octavalent dihedral oligomer^29^ (D4), and icosahedral cage^24^ (I53 or ICOS) (**Figure 1D and Table S1**). These structures fall into two broad geometric groups: (i) *cis*-DLL4 where DLL4 ligands are conjugated to one side of the structure (*i.e.*, C3-HB, C6, and C8), and (ii) *trans*-DLL4, where the ligands are affixed to opposite sides of the structure (*i.e.*, C2, C3, D3, D4, and ICOS) (**Figure 1C, D**). While C2 through D4 oligomers are homomeric proteins, the ICOS protein cage is a heteromeric protein complex assembled from the C3 and C5 proteins mentioned above, featuring 20 copies of C3 and 12 of C5, for a total of 120 single chains (**Figure 1D**).

**Figure 1:**
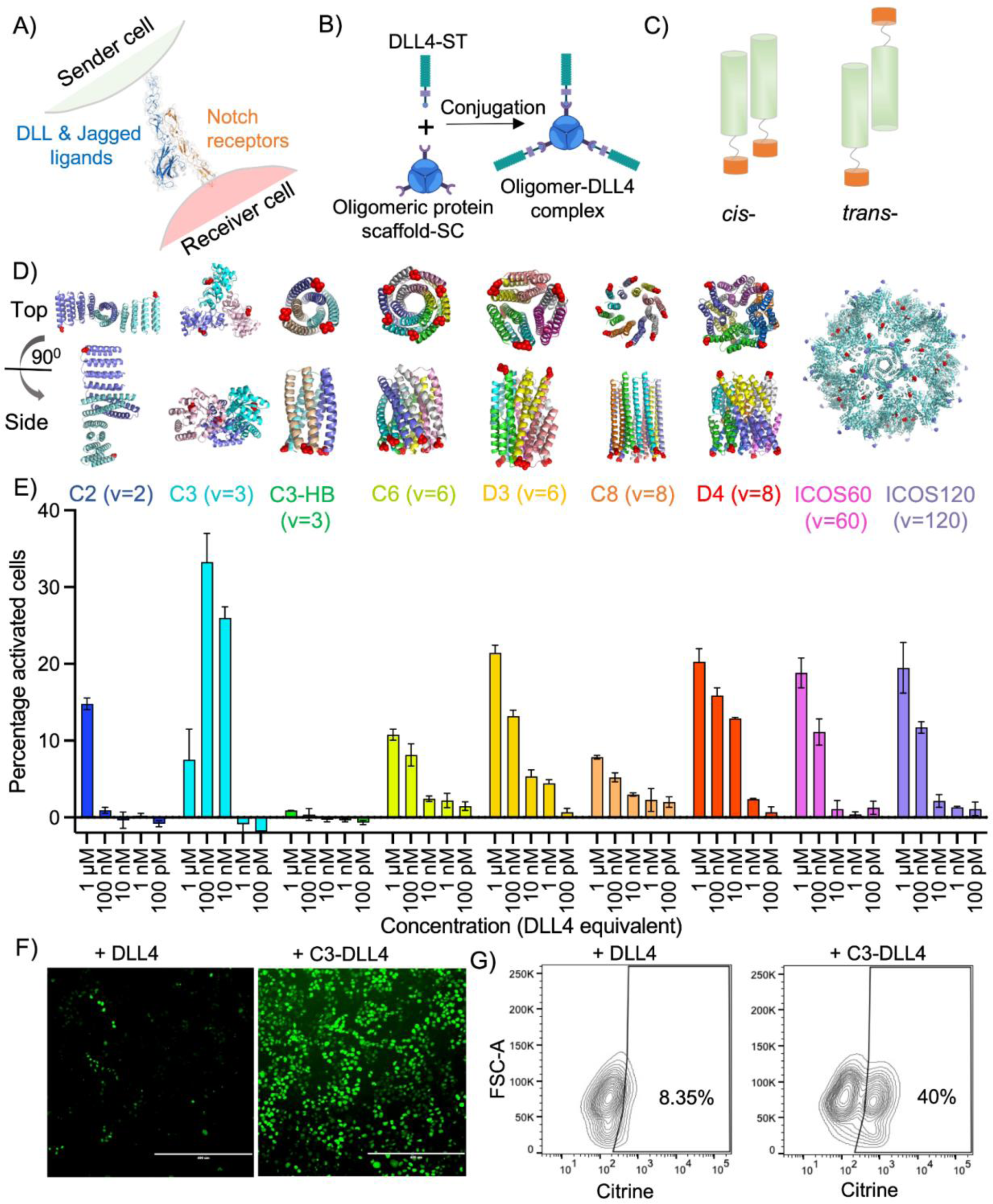
Engineering multivalent soluble Notch agonist. A) Schematic of Notch receptor activation by Delta-like ligands (DLL) or Jagged ligands. B) Conjugation of SpyTag Delta-like ligand 4 (DLL4-ST) with SpyCatcher (SC) oligomeric scaffolds to form a soluble multivalent DLL4 complex. C) Oligomeric scaffolds (green) presenting DLL4 (orange) in *cis*- or *trans*-configuration. D) PDB models of oligomeric protein scaffolds used in this study. C: cyclic; D: dihedral; ICOS: icosahedral; and V: valency. Red dots indicate the position of DLL4 conjugation. E) Flow cytometry showing the percentage of citrine-positive U2OS reporter cells for indicated concentrations of soluble Notch agonists. Concentrations reflect equimolar amounts of DLL4 ligand. F) Fluorescence microscopy, and G) flow cytometry data showing Notch activation in U2OS reporter cells upon 50 nM C3-DLL4 treatment. Scale bar = 400 µm.

Soluble DLL4 protein complexes were generated by mixing oligomer-SC with DLL4-ST at an equimolar ratio of DLL4 to the monomer of the oligomers (1:1 ratio of SC:ST). Consequently, the number of DLL4 ligands presented by these oligomeric protein complexes is equal to the state of multivalency: *i.e.*, C2-DLL4 = 2, C3-DLL4 = 3, C5-DLL4 = 5, C6-DLL4 = 6, D3-DLL4 = 6, C8-DLL4 = 8, D4-DLL4 = 8, ICOS60-DLL4 = 60, and ICOS120-DLL4 = 120. As the ICOS cage is assembled from 20 C3 and 12 C5 units, the total number of DLL4 ligands on the cage surface is either 60 (when 20 C3 OR 12 C5 units are conjugated to DLL4) or 120 (when 20 C3 AND 12 C5 are conjugated to DLL4). The resultant protein complexes were fully characterized by denaturing SDS-PAGE, native PAGE, and negative staining electron microscopy (EM) (**Figure S1**).

### Notch activation in an adherent reporter cell line

To assess Notch receptor activation by soluble oligomeric Notch ligands, we used an established Notch1-Gal4 reporter system that substitutes Gal4 in place of the NOTCH1 ankyrin repeat domain within the NICD to avoid interference from endogenous Notch signals^30,31^. Ligand binding and subsequent Notch receptor proteolytic processing releases a chimeric NICD1-Gal4 protein that can activate a citrine gene fused to Histone 2b (H2B) under the control of a Gal4-response element^31^. We assessed the degree of Notch activation by all oligomer-DLL4 complexes at multiple concentrations. At equimolar levels of DLL4 ligand, soluble C3-DLL4 exhibited the highest activation (**Figure 1E, F, G**), comparable to plate-bound DLL4; C2-DLL4, D3-DLL4, D4-DLL4, ICOS60-DLL4, and ICOS120-DLL4 induced intermediate Notch activation, and C3-SB-DLL4, C6-DLL4, and C8-DLL4 induced the lowest Notch activation (**Figure 1E**). We interpreted the differential activation of Notch signaling as a consequence of the structural and geometric differences in the oligomeric protein building blocks. C3-DLL4 has a flat structure with the DLL4 binding termini pointing outward in a plane (**Figure 1D, C3**), which we reasoned would facilitate the bridging of cells (*trans*-binding), thereby promoting Notch activation through cell-cell interactions and receptor clustering. Similarly, both D3-DLL4 and D4-DLL4 present ligands on opposite sides of the helical bundles, as do the icosahedral ICOS60-DLL4, and ICOS120-DLL4. However, while facilitating cell bridging, higher-order oligomeric structures likely present more unbound ligands, resulting in less efficient Notch activation than C3-DLL4. Those constructs showing the lowest degree of receptor activation, C3-SB-DLL4, C6-DLL4, and C8-DLL4 present ligands on one end of the long helical bundles, and thereby promote ligand binding to cells in *cis*, which would preclude bridging of cells. In addition, we tested an additional C3 complex (C3-1na0C3-DLL4) with a concave structure, which also presents ligands in *cis*. Predictably, this oligomeric-DLL4 complex likewise resulted in low levels of Notch activation (**Figure S2**).

### Structure-function relationship

We hypothesized that oligomeric DLL4 complexes with appropriate geometry, *i.e.*, *trans*-DLL4, might facilitate bridging between cells, and that Notch signaling might be activated through mechanical forces induced by cell movement. We incubated monomeric DLL4, dimeric C2-DLL4 and trimeric C3-DLL4 with U2OS-Notch1-Gal4 cells and assessed receptor binding. Immunostaining followed by confocal imaging showed that cells incubated with monomeric DLL4 failed to bind ligand within the concentration range tested, owing to low (micromolar) binding affinity to the Notch1 receptor^32,33^, and exhibited an even distribution of Notch1 receptors over the cell surface (**Figure 2A**). Cells incubated with C2-DLL4 showed DLL4 binding to the cell surface and co-localization with Notch1 in faint puncta; however, the Notch1 receptor remained evenly distributed on the cell surface (**Figure 2A** middle panel). In contrast, cells incubated with C3-DLL4 showed strong co-localization of the DLL4 ligand and the Notch1 receptor, with marked clustering at cell-cell junctions (**Figure 2A** bottom panel and **Figure S3**), indicating the formation of Notch synapses. Importantly, in C3-DLL4 treated cells, Notch1 was no longer evenly distributed on the cell surface and was only found at the C3-DLL4 mediated synapses. Notably, cells plated at low cell density and consequently lacking cell-cell contacts showed C3-DLL4 and Notch1 colocalization distributed evenly over the cell surface (**Figure S3A**). As these cells divided, C3-DLL4 and Notch1 re-localized to the cell-cell junction, forming synapses (**Figure S3B**). These synapses were observed for 5 days after C3-DLL4 treatment, indicating possible sustained Notch activation. Formation of synapses was also described recently in real-time imaging studies of endogenously tagged ligand and Notch receptor proteins in sender and receiver cells upon cell - cell contact^34^.

**Figure 2:**
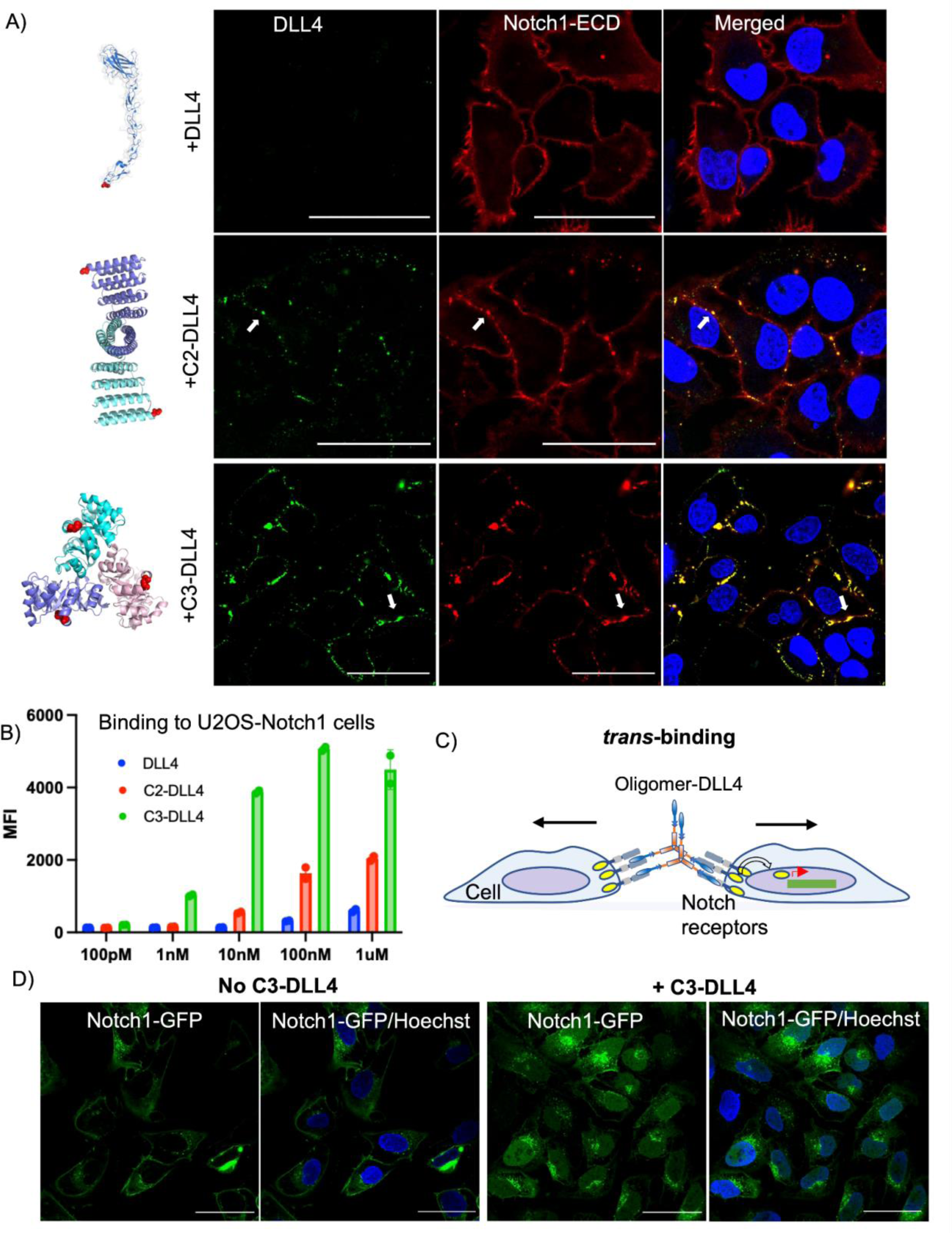
Mechanism of soluble agonist-mediated Notch receptor activation in U2OS reporter cell line. A) Confocal microscopy of U2OS-Notch1 cells after treatment with 10 nM equivalent of monomeric DLL4 (top panel), dimeric C2-DLL4 (middle panel), and trimeric C3-DLL4 (bottom panel). Immunostaining was performed for DLL4 (green), Notch1-ECD (red), and Hoechst (blue). The clustering of the Notch1 receptor and the co-localization of Notch1 and C3-DLL4 is marked by an arrow (middle and bottom panel). The binding of DLL4, C2-DLL4, and C3-DLL4 to U2OS-Notch1 cells was determined by immunostaining followed by flow cytometry. C) Models showing *trans*-binding of oligomeric DLL4 which results in Notch1 receptor clustering and synapse formation. D) The release of Notch intracellular domain NICD1-GFP induced by C3-DLL4. Confocal microscopy of U2OS-Notch1-GFP cells in the absence (left panel), or presence (right panel) of C3-DLL4 (50 nM), showing NICD1-GFP on the membrane (no activation, top panel), or in the nucleus (24 h after activation, bottom panel), with Hoechst (blue) nuclear staining. Scale bar = 50 µm.

The ability to cluster Notch1 receptors at cell-cell junctions is enhanced by the cooperative binding activity of the oligomeric-DLL4. Immunostaining followed by flow cytometry showed that trimeric C3-DLL4 showed the highest binding to U2OS-Notch1 cells; dimeric C2-DLL4 showed intermediate binding, whereas monomeric DLL4 exhibited the lowest binding (**Figure 2B**). Collectively, both the geometry (*i.e.*, *trans*-binding capability) and multivalency (binding affinity) of the oligomeric-DLL4 appear to contribute to receptor clustering and Notch synapse formation (**Figure 2C**), which in turn dictate signal activation, with C3-DLL4 eliciting higher activation than C2-DLL4 (**Figure 1E**).

To track NICD1 translocation to the nucleus, we used a cell line expressing a Notch1 receptor in which Notch1 was fused to GFP at its C-terminus (U2OS-Notch1-GFP). NICD1-GFP translocated to the nucleus after C3-DLL4-mediated Notch activation, whereas cells lacking exposure to C3-DLL4 exhibited Notch1 fluorescence at the cell surface only (**Figure 2D**). In addition, cells seeded at low density to avoid cell-cell contact did not show nuclear translocation of NICD1-GFP, indicating a requirement for Notch synapse formation for signal activation (**Figure S4).** Time-lapse live cell imaging revealed that visible accumulation of NICD1-GFP in the nucleus took between 3-10 hours (**Figure S5**). To demonstrate downstream Notch activation following nuclear translocation of NICD1, we used a reporter cell line engineered to express a 12xCSL - mCherry target. 48 hours after C3-DLL4-mediated Notch activation, we detected both mCherry and nuclear NICD1-GFP (**Figure S6**).

### Notch activation in a suspension reporter cell line

To assess Notch activation in suspension culture, we engineered K562 cells to express a Doxycycline (Dox)-inducible Notch1-Gal4 chimera as above, and a UAS-mCherry reporter (K562-Notch1-Gal4 cells). The Notch1-Gal4 chimeric protein also contains a FLAG tag at the N-terminus to facilitate detection of the Notch1 extracellular domain. Addition of C3-DLL4 at different concentrations induced cell clustering. At high concentrations of C3-DLL4 (1 µM), cells remained homogeneously dispersed; however, at lower ligand concentrations, cells formed clusters, most prominently at 10 nM (**Figure 3A**). As the lowest concentration of C3-DLL4, cells were homogeneously distributed, as were control cells that were not exposed to ligand (**Figure 3A**). In contrast, we observed absent or minimal clustering for soluble C5-DLL4, C6-DLL4, and C8-DLL4. Further, ICOS60/120-DLL4 constructs clustered cells in a concentration-dependent manner, similar to C3-DLL4 (**Figure S7**).

**Figure 3:**
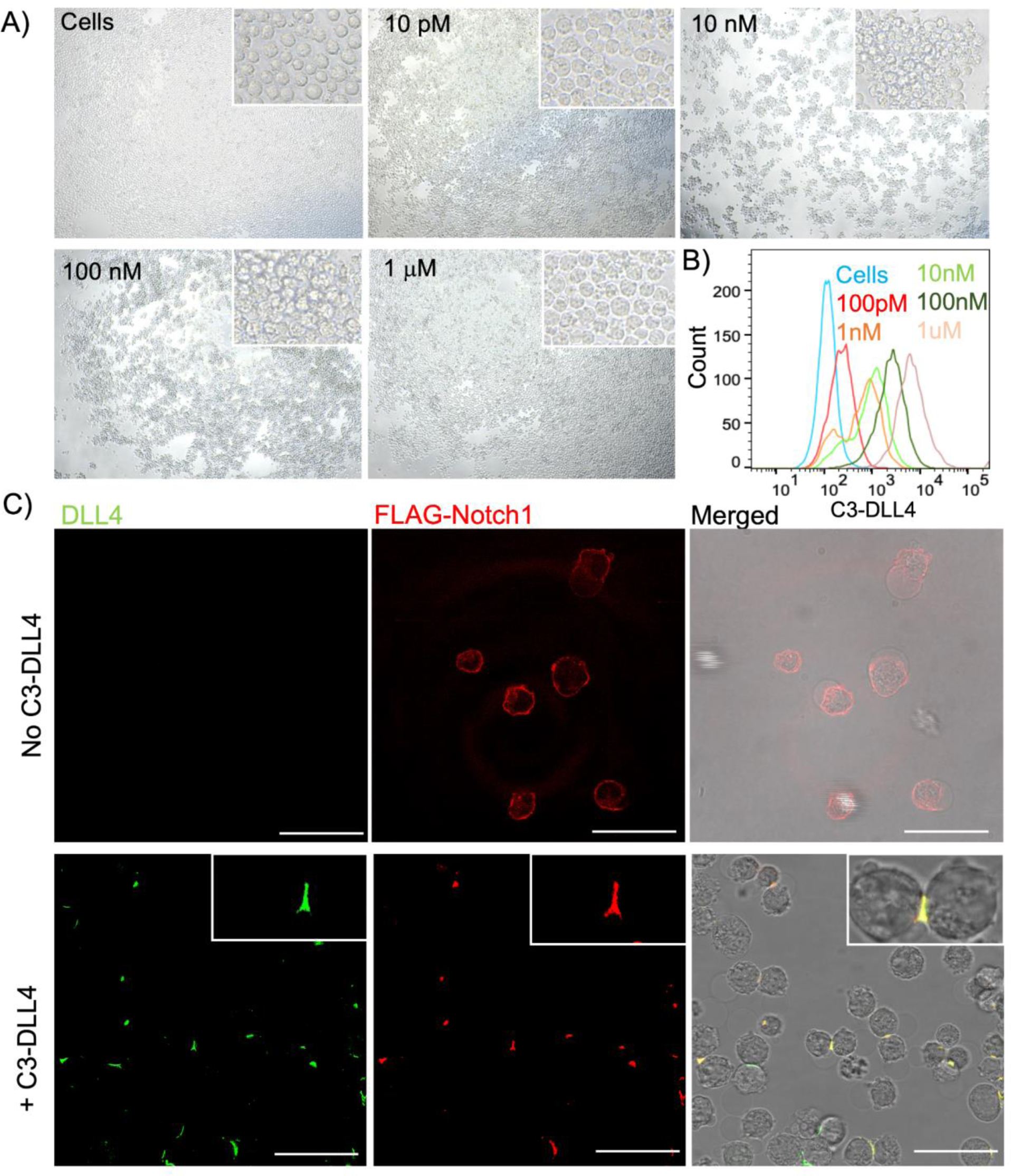
Multivalent Notch agonists induce clustering in suspension cells. A) Clustering of K562-Notch1-Gal4 cells induced by C3-DLL4 complex in a ligand concentration-dependent manner. Phase contrast micrograph presenting clustered cells at various C3-DLL4 concentrations. Insets: higher magnification showing cellular clusters. B) Flow cytometry data showing the binding of C3-DLL4 to K562-Notch1-Gal4 cells at various ligand concentrations, where cell-bound C3-DLL4 was stained with Anti-His antibody. The intensity indicates the degree of C3-DLL4 binding to the cell surface. C) Notch synapse formation through cell-cell interaction in the absence (top panel) or presence (bottom panel) of C3-DLL4 (10 nM). Immunostaining was performed for DLL4 (green), and Notch1-ECD (using a FLAG epitope at the N-terminal end of the Notch1 ECD) (red). Inset showing C3-DLL4/Notch1-ECD colocalization through receptor clustering and formation of a Notch synapse (bottom panel). Scale bar = 50 µm.

We sought to investigate the mechanism of the concentration-dependent cell-cell interaction and subsequent clustering induced by C3-DLL4. We hypothesized that at high concentrations, C3-DLL4 binding to Notch receptors on the K562 reporter cells reaches saturation, and thus precludes cell bridging—a phenomenon akin to the Hook effect (**Figure S8**)^35^. Lower concentrations of C3-DLL4 leave unbound Notch receptors on the cell surface to interact with multivalent DLL4 ligands bound to other cells, thereby facilitating cell bridging ( **Figure S8**). To test this hypothesis, we incubated the K562 reporter cells with C3-DLL4 at various concentrations for 1 hour. Before cluster formation, we assessed the binding of C3-DLL4 on the cell surface by immunostaining and flow cytometry. Cells stained at 1 µM C3-DLL4 showed the highest fluorescence intensity (**Figure 3B**). As the concentration of C3-DLL4 decreased, the fluorescence intensity of the cells decreased; at a concentration of 10 nM, cells were segregated into a major C3-DLL4-positive and a minor C3-DLL4-negative population (**Figure 3B**). Moreover, the ligand-bound population had lower mean fluorescence intensity compared to cells incubated at 1 µM concentration, indicating that lower C3-DLL4 concentrations did not fully saturate Notch receptor binding. Under non-saturating conditions, free Notch1 receptors on one cell can interact with an unoccupied ligand of C3-DLL4 bound to a neighboring cell (**Figure 3B**). Taken together, these data establish that C3-DLL4 can bridge cells to induce cell-cell interaction between K562 reporter cells at a suitable concentration, forming cellular clusters.

### Notch synapse in suspension cells

Clustered K562 reporter cells were fixed and stained for C3-DLL4 and for the ECD of Notch1. Both C3-DLL4 and Notch1 were localized at the cell-cell junctions, consistent with Notch synapse formation, as observed recently (**Figure 3C**, bottom panel; **Figure S9**)^34^. Untreated control cells showed a more even surface distribution of Notch1 (**Figure 3C**), while cells treated with C3-DLL4 for 1 hour, before clustering was detected, showed bound C3-DLL4 distributed more evenly on the cell surface, co-localized with Notch1 (**Figure S9**). Clustered cells could be dissociated by applying a rotational force (orbital shaker, 100 rpm) to induce the expression of the reporter gene (mCherry) following Notch synapse formation (**Figure S10**).

### Endogenous Notch activation

Next, we tested whether the multivalent soluble Notch agonists could activate signaling from endogenous Notch receptors. We tested MDA-MB-231, a triple negative human breast cancer cell line, and SVG-A, a human fetal astrocyte cell line, both of which express endogenous Notch. To synchronize Notch activation, we incubated C3-DLL4 with the cells in the presence of 500 nM γ-secretase inhibitor (GSI, compound E) for 24 hours and removed the GSI by media replacement. Three hours after media replacement, we performed RT-qPCR for Notch target genes Hes1, Hey1, Hey2, HeyL, and NRARP in MDA-MB-231 cells, and for Hes1, Jag1, TRIB1, Hey1, Hey2, and HeyL in SVG-A cells (**Figure S11)**, and observed upregulation. Western blot analysis showed receptor cleavage to generate NICD1 (**Figure S11**), confirming the activation of endogenous Notch1 by C3-DLL4.

### Soluble Notch agonists support in vitro T cell differentiation

Notch signaling is indispensable for T cell development from hematopoietic stem and progenitor cells (HSPCs). Because of the biomechanical mechanisms required for Notch receptor activation, current protocols for *in vitro* T cell differentiation typically present Notch ligands coated on culture plates or expressed by feeder cells^15,17,36,37^. To test whether soluble multivalent DLL4 complexes can support *in vitro* T cell differentiation similarly to the immobilized Notch ligands, we treated human cord blood (CB) derived CD5^+^CD7^+^ T cell progenitors with our designed series of oligomer-DLL4 complexes during a two-week induced differentiation into CD4^+^CD8^+^ double positive (DP) T cells^16^. Confirming the optimal geometry and valency within our series, C3-DLL4, but not other multivalent DLL4 complexes, supported the derivation of CD4^+^CD8^+^ DP T cells (**Figure 4A, B**). These results indicate that soluble C3-DLL4 activates the Notch signal in both engineered reporter systems and in a more biologically relevant condition that mimics T cell development. Additionally, when C3-DLL4 was added during the early stage of T cell differentiation, we generated lymphoid progenitors (CD7^+^) and T cell progenitors (CD5^+^CD7^+^) from iPSC-derived HSPCs (**Figure 4C**), further suggesting that the C3-DLL4 supports Notch activation at different stages of T cell differentiation, and is not only restricted to CB-derived cells.

**Figure 4:**
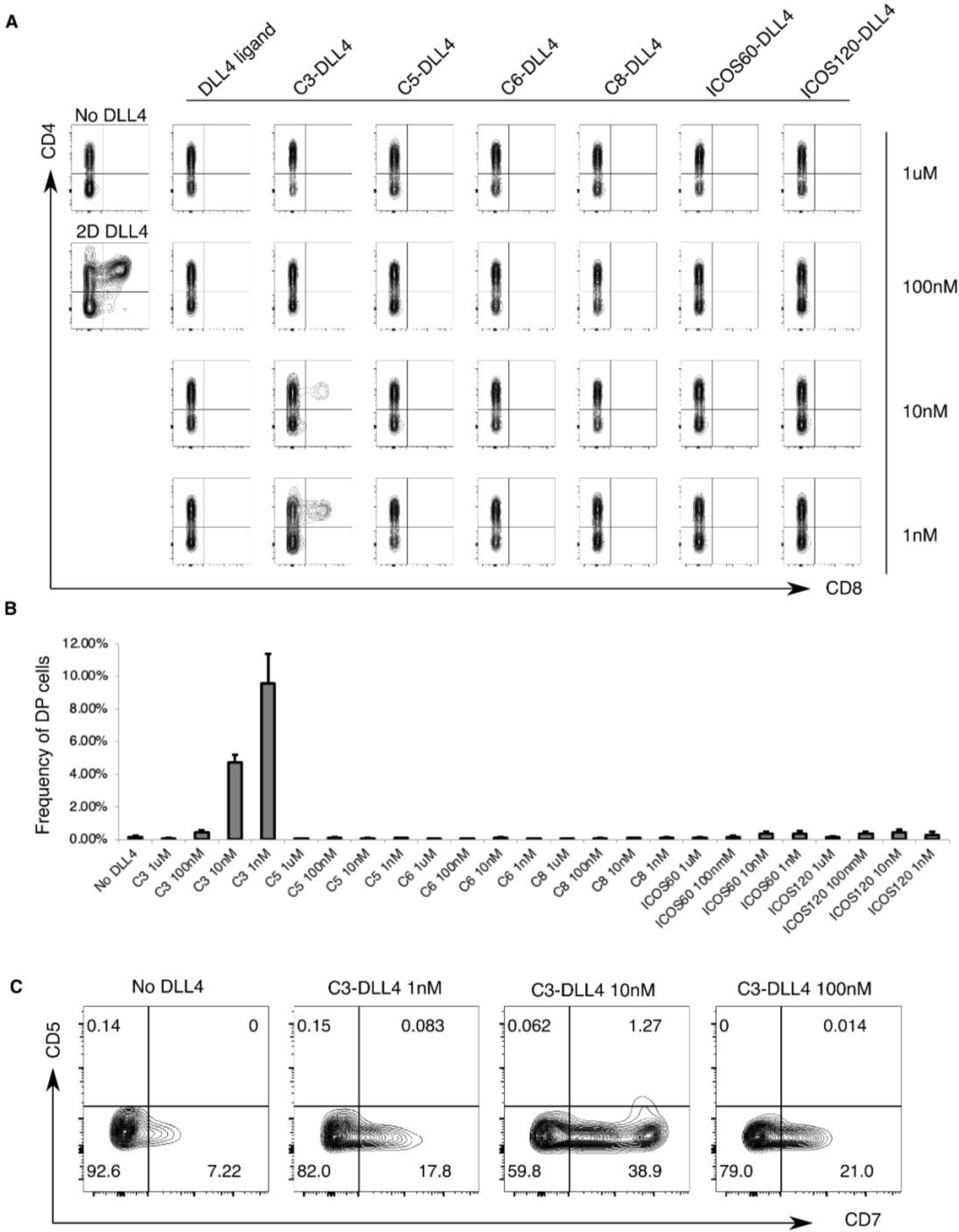
Multivalent Notch agonists support *in vitro* T cell differentiation. A) Flow cytometry showing expression of CD4 and CD8 in CB-derived proT cells cultured with no DLL4, 2D-DLL4 (10 µg mL^-1^), or oligomer-DLL4 complexes during two weeks of double positive (DP) T cell induction. B) Bar chart showing frequencies of DP T cells after the treatment with oligomer-DLL4 complexes. C) Flow cytometry shows the expression of CD7 and CD5 in iPSC-derived HECs cultured with no DLL4 or C3-DLL4 complexes during two weeks of proT cell induction.

Having shown that soluble C3-DLL4 can support *in vitro* T cell differentiation, we next explored if the soluble C3-DLL4 can facilitate the derivation of T cells from iPSCs in 3D bioreactors, thereby enabling large-scale manufacturing of cell therapy products^38^. Human iPSCs were induced to form embryoid bodies, dissociated into adherent hemogenic endothelial cells (HECs), and cultured during two weeks of lymphoid specification to give rise to lymphoid/T progenitors^16^. From floating cells, we collected CD5^+^CD7^+^ T cell progenitors and differentiated the cells into T cells via an established 2D culture condition using immobilized DLL4, or in a bioreactor for suspension culture with C3-DLL4 (**Figure 5A**). After two weeks of differentiation, CD4^+^CD8^+^ DP T cells were produced in both 2D and 3D conditions. Mature CD3^+^TCRαβ^+^ T cells were detected in bioreactor cultures (**Figure 5B**). Importantly, although bioreactor cultures produced T cells at a lower cell density than 2D cultures, bioreactor cultures yielded a significantly greater overall number of CD3^+^TCRαβ^+^T cells per volume of culture media, and an increased number of T cells derived per microgram of DLL4 ligands (**Figure 5C, D**). These data establish that soluble C3-DLL4 supports a more scalable, cost-efficient platform for iPSC-T cell production by enabling T cell derivation in 3D suspension culture.

**Figure 5:**
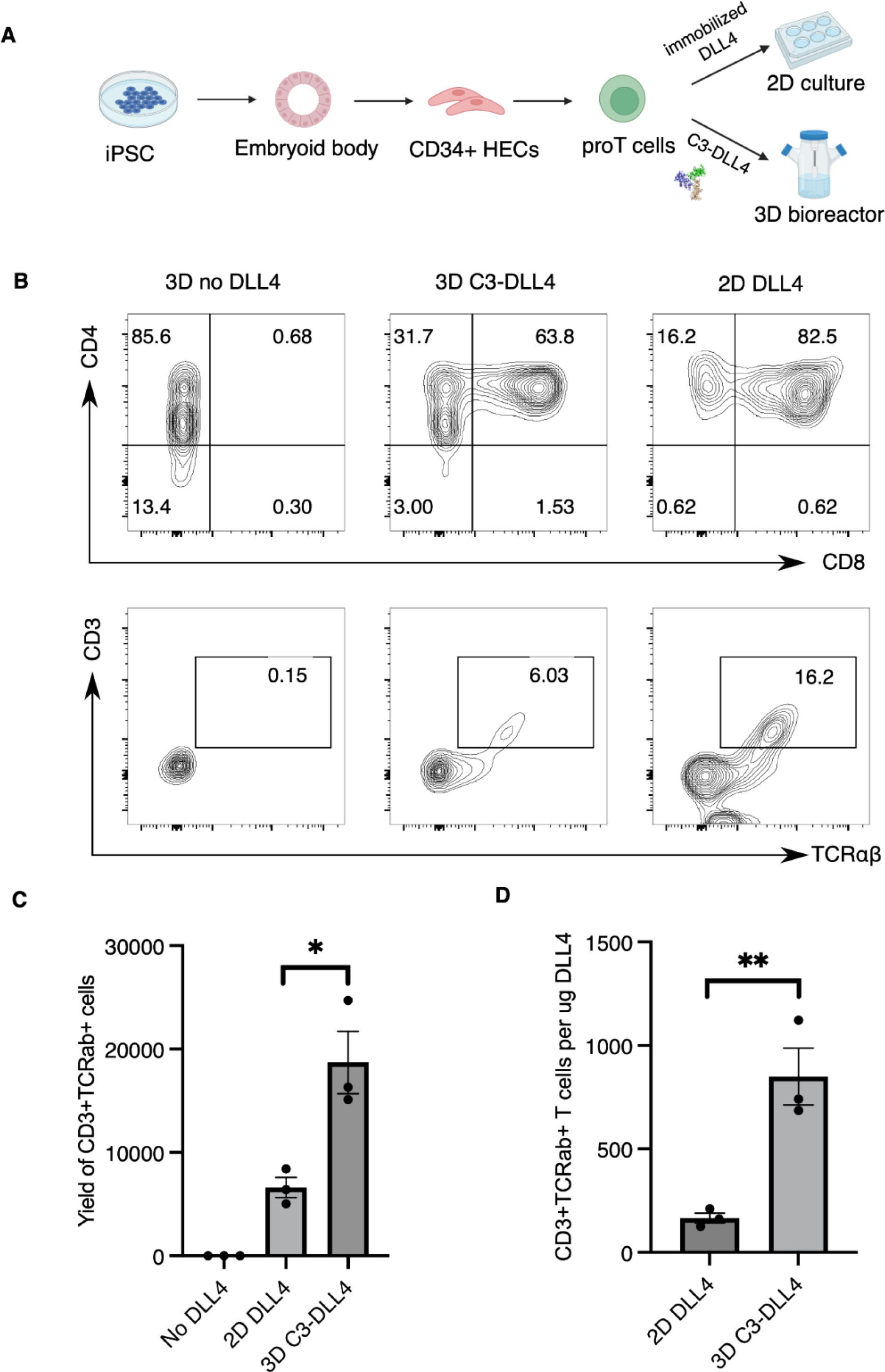
Multivalent Notch agonist facilitates derivation of iPSC-T cells via 3D bioreactor culture. A) Schematic illustration of experimental design. B) Flow cytometry results showing expression of T cell markers (CD4, CD8, CD3, TCRαβ) in iPSC-derived T cells (week 4) cultured with 2D-DLL4 (10 µg mL^-1^) or 3D bioreactor (No DLL4, 10 nM C3-DLL4), C) Bar chart showing yield of CD3^+^TCRαβ^+^ iPSC-T cells (week 4) derived via 2D-DLL4 (10 µg mL^-1^) or 3D bioreactor (No DLL4, 10 nM C3-DLL4) from 400K proT cells. D) Bar chart showing the number of CD3^+^TCRαβ^+^ iPSC-T cells (week 4) derived using 1 µg of DLL4 via either a 2D or 3D bioreactor culture system.

## Conclusion

Optimizing the amplitude and temporal control of receptor signaling through protein engineering serves numerous applications in biomedicine. Leveraging synthetic biology, *de novo*-designed protein scaffolds and receptor binders have been developed to control cytokine signaling in hematopoiesis^20–22^, providing precise control over ligand presentation through appropriate geometries and valencies to fine-tune the signal intensity. In addition, modulating cell-cell interactions through extracellular ligand engineering has recently emerged as a new paradigm for controlling multicellular organization and signal transduction^39,40^. Within the field of immunotherapy, agents that induce heterotypic cell-cell interactions, including bispecific antibodies (bispecific T cell engagers, BiTEs), have been clinically approved to treat various cancers^41^.

Notch signaling is central to many cellular differentiation processes, most notably T cell development, where tools to activate Notch signaling in a regulated, precise manner would facilitate *in vivo* strategies and 3D bioreactor manufacturing. Designing a soluble cytokine-like Notch agonist, however, presents specific challenges owing to the nature of Notch signaling in which cell-bound transmembrane ligands deliver mechanical force to activate receptors on juxtaposed cells. There are no endogenous soluble ligands, and free soluble Notch ligands inhibit Notch signaling^6^. By screening a panel of multivalent Notch ligands with different geometries and ligand presentation, we identified an optimal multivalent configuration that bridges cells and promotes receptor clustering. The bridging of suspension cells by C3-DLL4 – Notch interactions provides the tethering necessary for the delivery of the mechanical force that activates Notch receptor signaling, thereby enabling cell amplification in liquid suspension cultures and delivery of Notch ligands *in vivo* to develop immunotherapy. Consistent with the interpretation that mechanical shear experienced by Notch proteins on ligand-bridged cells in suspension culture is sufficient to induce the conformational shifts that favor receptor activation, we found that ligand-mediated Notch receptor clustering preceded signal activation. The geometry of the designed scaffold protein and the orientation of ligands appears critical: Designed helical bundles that presented Notch ligand in a trimeric *trans*-binding configuration best promote Notch synapse formation, with receptor clustering at the cell-cell junction presumably providing a ‘signaling hub’ for amplified Notch activation. A recent study has shown that Notch activation can occur through Notch synapse formation at sender-receiver cell junctions, consistent with our C3-DLL4 mediated receptor clustering^34^.

We have demonstrated the utility of C3-DLL4, a trimeric Notch agonist, in T cell generation from cord blood and human iPSCs, and provide a proof-of-principle for C3-DLL4 mediated T cell production in a 3D bioreactor. Of note, in comparison to previous DLL microbead-based Notch activation for T cell generation^14^, our method provides a more precise way to induce Notch activation through the appropriate control of the agonist characteristics such as a well-defined geometry, valency, and homogeneity. Moreover, the use of protein-based soluble Notch agonists may provide an avenue to control Notch activity in *in vivo* systems. Other Notch ligands (*i.e.*, DLL1, JAG1, and JAG2) and ligands that activate other signaling systems could easily be integrated into this 3D suspension culture system to further optimize T cell production. We envision that innovation in using designed protein tools for 3D bioreactor production of therapeutic T cells will significantly improve yield and lower production costs.

Beyond T cells, a large variety of cellular systems require Notch activation, including the generation of definitive HSCs and dendritic cells (DCs)^42^. Since HSCs show promise for bone marrow transplantation and DCs for cancer vaccines, we envision that our synthetic Notch agonists will be useful tools for the commercial manufacturing of these cells. Likewise, neurons, vascular endothelial cells, and other cell types require Notch signaling as a fate specifier. Many of these cells differentially express Notch receptors and require varying amplitudes of signal activation. Often these cells are grown in 3D organoid culture, where 2D Notch activation is not feasible. Soluble cytokine-like Notch agonists, which would penetrate more deeply into different layers of an organoid, could be valuable adjuncts to existing toolkits for organoid production, manipulation, and investigation. Thus, our method could be further extended to build designer Notch agonists that will suit a particular cell type and differentiation condition for precise signal activation.

